# Non-linear tumor-immune interactions arising from spatial metabolic heterogeneity

**DOI:** 10.1101/038273

**Authors:** Mark Robertson-Tessi, Robert J. Gillies, Robert A. Gatenby, Alexander R. A. Anderson

**Author notes:** M. Robertson-Tessi, R. J. Gillies, R. A. Gatenby and A. R. A. Anderson are with the Moffitt Cancer Center, Tampa, FL 33612 USA.

## Abstract

A hybrid multiscale mathematical model of tumor growth is used to investigate how tumoral and microenvironmental heterogeneity affect the response of the immune system. The model includes vascular dynamics and evolution of metabolic tumor phenotypes. Cytotoxic T cells are simulated, and their effect on tumor growth is shown to be dependent on the structure of the microenvironment and the distribution of tumor phenotypes. Importantly, no single immune strategy is best at all stages of tumor growth.

## I. Tumor-Immune Interactions

Both the innate and adaptive arms of the immune system have been shown to induce tumor cell death [1, 2]. Immune surveillance is widely thought to eradicate many microscopic cancers before they become clinically-apparent. Conversely, clinical cancers are assumed to have developed adaptive strategies to evade immune attack. We examine a simplified tumor-immune model that pits T cells against a small growing tumor. In some cases, the immune system is able to remove a small tumor before it becomes detectable, but here we focus on the case where a tumor does eventually escape, and look at the tumor-immune interactions. There are multiple mechanisms of immune evasion that tumors use, including immunosuppressive surface markers such as PD-L1 [3], down-regulation of antigen presentation machinery [4], recruitment of immunosuppressive immune cells [5], and secretion of immunosuppressive factors such as TGF-β [6] and extracellular acid [7]. We focus on the latter here, a consequence of the altered glycolytic metabolism seen in many tumors.

## II. Tumor-Environment Heterogeneity

Heterogeneity is increasingly being recognized as an important aspect of tumor biology. Recently, Swanton and colleagues [8] showed that multiple biopsies from the same tumor display distinct genetic profiles and yet are phenotypically similar. This phenotypic convergence despite genotypic divergence has previously been examined theoretically [9] and may be a predictable evolutionary consequence of the tumor ecosystem [10, 11]. Quantifying this heterogeneity and the dynamics of its evolution remain a challenge, as does the understanding of how it relates to overall outcome [12–14]. Here we examine an environment that is temporally and spatially heterogeneous, largely due to variations in blood flow, resulting in local fluctuations of nutrients as well as variable infiltration of immune cells. The tumor cells are allowed to evolve, and selection pressures in each niche lead to the development of different tumor cell phenotypes. This heterogeneity then has implications for the immune response, both with respect to the tumor cells’ properties and the environments in which they reside.

## III. Results

Using a two-dimensional hybrid cellular automaton model, we simulated growing tumors in a vascularized tissue [15]. Heterogeneity was incorporated on two metabolic axes: glycolytic capacity and resistance to extracellular acidosis. Absent treatment, a subset of tumor cells evolve under environmental selection pressure to become glycolytic and acid resistant. These properties cause increased tumor invasiveness under the right heterogeneous conditions.

Within this context, immune pressure in the form of migratory cytotoxic T cells was added to the model. Figure 1 shows the results of tumors growing under three different antigenic values, compared to a simulation with no T cells. The antigenicity is simply modeled as the rate at which T cells enter the domain per unit time, given a particular tumor size. The tumor has two growth phases: 1) the initial, slow growing phase when the metabolic phenotypes are still benign, and 2) the invasive phase, in which acid-mediated invasion drives fast growth. of note is the differential effects of the immune response on these two phases, with the switch occurring near the ‘elbow’. In the first part of the growth, increased antigenicity leads to slower growth rates for the tumor, expected since more T cells equates with faster tumor cell killing and therefore checked growth. However, near the elbow and beyond, it is clear that increases in antigenicity have a non-linear effect, in that the highest value leads to faster development of an invasive tumor than a moderate antigenicity. This can be understood in the context of spatial heterogeneity. The poor vasculature near the center of the tumor promotes acidosis, as well as limiting the intravasation of T cells; this leads to the immune response being primarily applied on the edges of the tumor. However, the most aggressive metabolic phenotypes develop in the center, and this differential killing allows them to escape and become invasive sooner. In other words, the immune system, if too aggressive, can destroy the spatial heterogeneity that was keeping invasive cell phenotypes in a dormant state.

**Figure 1.**
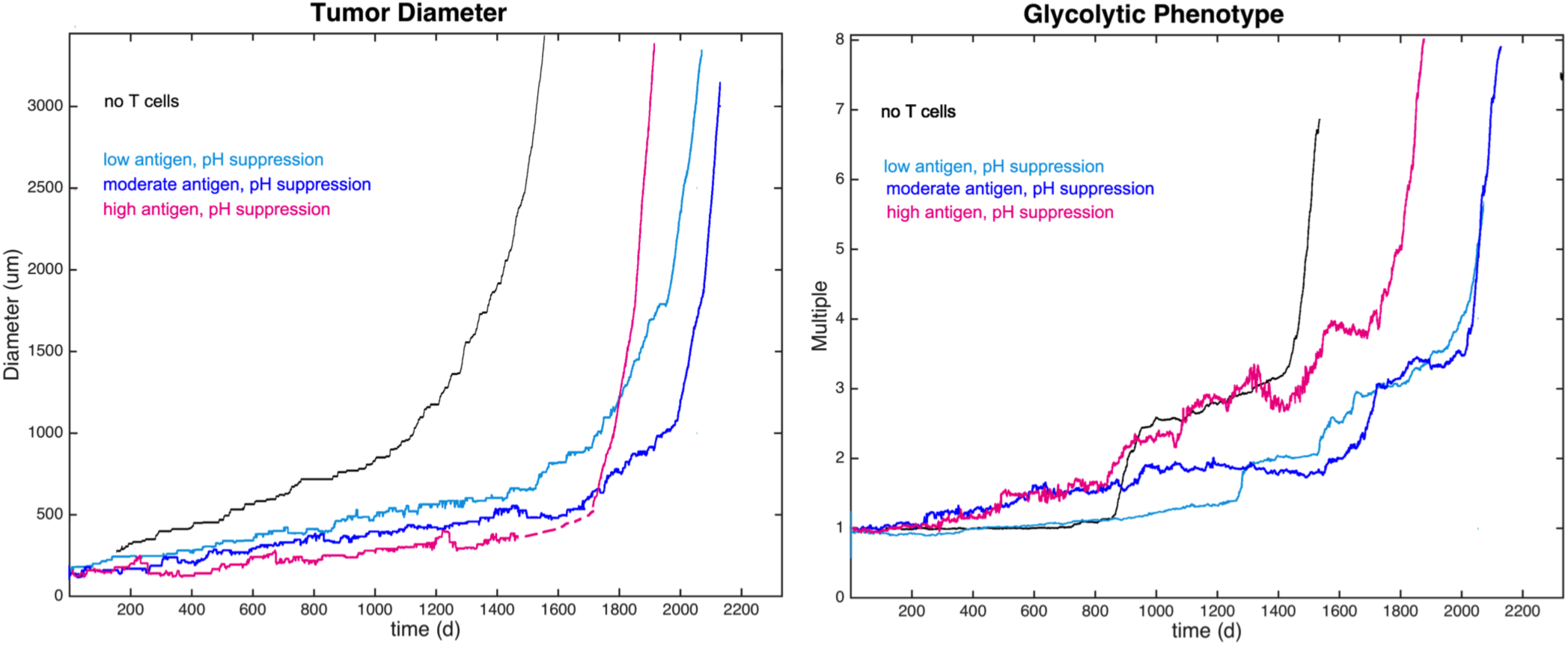
Left: Simulations with no T-cells and three levels of antigenicity (see colors in legend). Increasing antigenicity slows tumor growth in early phase (t<1500). Near the ‘elbow’ (dashed pink), the highest antigenicity leads to faster tumor invasion compared to low antigenicities. Right: Glycolytic phenotypes are selected by medium and high antigenicities early on, despite slower tumor growth, leading to faster emergence of invasive cells.

## IV. Quick Guide to the Methods

### A. Equations

The concentration of a molecule (C(x)) across a tissue is described by

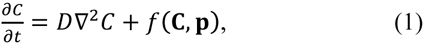

with diffusion constant *D*, and *f* describing the production and consumption of the molecule depending on the concentrations of extracellular molecules (**C**(*x*)) and cellular parameters (**p**(*x*)) at position *x*. Cells primarily produce ATP from glucose (*G*), using either an efficient aerobic pathway that requires oxygen (*O*), or using glycolysis, an inefficient anaerobic pathway that produces protons (*H*). The model assumes that cells meet a target level of ATP demand by preferentially using the aerobic pathway, and making up the difference by increasing flux through the glycolytic pathway in hypoxic regions. Oxygen consumption (*f*_*O*_) and glucose consumption (*f*_*G*_) are determined by the need to meet normal ATP demand (A_0_), given by

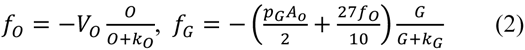

For tumor cells, if the coefficient *p*_*G*_>1, the tumor will consume more glucose than needed to meet normal ATP demand, representing constitutively activated glucose consumption seen in many tumors. The actual ATP production rate for the cell (*f*_*A*_) is determined from nutrient consumption rates, given by

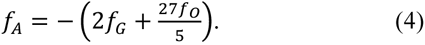

Proton production (*f*_*H*_) is linked to the amount of glycolysis that does not feed the aerobic pathway, given by

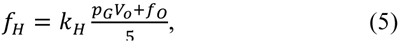

where parameter *k*_*H*_ accounts for proton buffering.

This metabolic program is implemented into each cell of a hybrid cellular automaton (HCA) model. One cell type is permitted per grid point, either a normal cell, tumor cell, necrotic cell, or blood vessel. For each time step *dt*, Eq. (1) is solved over the domain of the HCA for the steady state. Then, cells in the grid are put through a decision process based on the metabolic state of each cell and the nutrient concentrations at that point. Cells that have enough ATP production to meet the threshold of proliferation (*A*_*q*_) will advance their cell cycle. Cells that have completed the cell cycle will proliferate if there is adjacent space. The cycle is not advanced if the cell is quiescent due to lowered ATP production. Cells with production less than a death threshold are removed.

Tumor cells in the model have two continuously variable, heritable traits: excess glucose consumption, *p*_*G*_ from Eq. 3; and resistance to extracellular acidosis. These traits are passed from a parent to its two daughter cells with some small variation, chosen at random from an interval equally weighted in both directions to avoid biased drift.

A point-source vasculature is used to simulate blood vessels that spatiotemporally deliver nutrients and remove waste products. The field of vessels is seeded using a circle-packing algorithm based on vessel densities *in vivo*. This initial distribution can be altered by the creation of new vessels through angiogenesis, or by vessel degradation. For angiogenesis, new vessels are added to regions of hypoxia until there is enough oxygen delivery to remove the hypoxic state. Vessels are degraded over time due to surrounding tumor growth until they are lost from the tissue. These two opposing vascular forces impact the gradients of diffusible molecules.

T cells are modeled as individual cells on a separate grid, added to the layer with a rate (the ‘antigenicity’) that is proportional to the tumor size. T cells are added at any vessel that is within 100 microns of the tumor; once in the field, they migrate randomly. Upon encountering a tumor cell, they have a probability of killing it; said probability is lowered in acidosis. After a fixed number of kills, the T cell expires and is removed from the layer.

### B. Type of settings in which these methods are useful

HCA models have been used extensively to model cancer, including applications to angiogenesis [16, 17], cell motility and invasion [18, 19], tumor-immune interactions [20], and metastasis [21, 22].

